# An Anatomically and Hemodynamically Realistic Simulation Framework for 3D Ultrasound Localization Microscopy

**DOI:** 10.1101/2021.10.08.463259

**Authors:** Hatim Belgharbi, Jonathan Porée, Rafat Damseh, Vincent Perrot, Léo Milecki, Patrick Delafontaine-Martel, Frédéric Lesage, Jean Provost

## Abstract

The resolution of 3D Ultrasound Localization Microscopy (ULM) is determined by acquisition parameters such as frequency and transducer geometry but also by microbubble (MB) concentration, which is also linked to the total acquisition time needed to sample the vascular tree at different scales. In this study, we introduce a novel 3D anatomically- and physiologically-realistic ULM simulation framework based on two-photon microscopy (2PM) and in-vivo MB perfusion dynamics. As a proof of concept, using metrics such as MB localization error, MB count and network filling, we could quantify the effect of MB concentration and PSF volume by varying probe transmit frequency (3-15 MHz). We find that while low frequencies can achieve sub-wavelength resolution as predicted by theory, they are also associated with prolonged acquisition times to map smaller vessels, thus limiting effective resolution. A linear relationship was found between maximal MB concentration and inverse point spread function (PSF) volume. Since inverse PSF volume roughly scales cubically with frequency, the reconstruction of the equivalent of 10 minutes at 15 MHz would require hours at 3 MHz. We expect that these findings can be leveraged to achieve effective reconstruction and serve as a guide for choosing optimal MB concentrations in ULM.

## I. INTRODUCTION

For long, we were bound by the limit of diffraction stated by Rayleigh’s criterion [1] in conventional ultrasound imaging. Diffraction causes the point-spread function (PSF) to have a specific size in the order of the wavelength, which increases with the penetration depth: larger depths can be achieved at the cost of larger wavelength and thus degraded resolution. Wavelength thus dictates the compromise between limiting factors of resolution and penetration depth in conventional ultrasound [2]. The concept of isolated source localization from optical imaging [3], used in ultrasound localization microscopy (ULM) [2], [4], [5] challenges this compromise.

Microbubbles (MB) injected into the bloodstream are used as small, highly echogenic unique scatterers that can be tracked through vascular networks in large vessels down to, in principle, the smallest capillaries. It is by finding the exact centroid of each MB in reconstructed images or by fitting a paraboloid in RF data that we can obtain micrometer-scale resolution. This resolution is no longer dependent on wavelength, but rather on arrival time variance, number of elements, array apertures, distance between the array and MB, and sound speed [6].

The gain in spatial resolution comes at the cost of temporal resolution (in the order of a few minutes depending on the vascular network density) since sampling the vascular network requires localization and tracking of millions of MB in tens of thousands of images [7].

In [2], Couture et al. defined four conditions that are required to perform ULM, namely 1) being sensitive to the contrast agent, 2) being able to localize that contrast agent, 3) record at a high framerate and 4) maintain isolated sources in a spatiotemporal referential. our ability to respect the latter condition depends mainly on two factors: 1) acquisition parameters such as frequency and transducer geometry [6], and 2) microbubble concentration [7].

To quantify ULM limitations, studies have been performed in in-vitro phantoms with various materials, configurations, and scale. Viessmann et al. [8] conducted four experiments at 2-2.5 MHz with the last one using two touching cellulose tubes with inner diameters (ID) of 200 μm. Two sets of MB could be distinguished with a subdiffraction limit with a measured center-to-center distance of 197 μm (wavelength of 616-770 μm), but this setup was limited to relatively large mono-diameter tubes. Similarly, in [9], O’Reilly and Hynynen conducted an experiment at 612 kHz using a PTFE tube with 255 μm of ID spiraled around a 2.5 mm rod. The estimated localization uncertainty was again sub-diffraction limit with approximately 1/120 and 1/60 of the wavelength, but the limiting factor was the size of the phantom. In [10], Couture et al. succeeded in localizing MB in 100 μm sized microfluidic systems at 5 MHz. Similarly, in [11], Desailly et al. succeeded in localizing MB in rectangular 40-100 μm by 80 μm PDMS channels out of printed SLA molds separated by 50 to 200 μm at 1.75 MHz. However, none of the above setups’ scale, complexity and realism reflects *in-vivo* challenges. In mice, brain vascular networks typically exhibit large vessels with diameters in the 50-200 μm range [12] down to very small vessels in the <10-micrometer range [13]. As for how close vessels are to each other, vascular density varies from 161-391 sections per mm^2^, or 51-79 micrometers between them [14]. Furthermore, the size of the vessel intrinsically dictates the number of flowing MB and their velocities, which affects effective resolution [7]. To our knowledge, there is no validation framework for ULM image formation algorithms based on anatomically-realistic MB flow to assess ULM limitations.

To overcome this problem, a highly resolved ground truth of brain vasculature is needed. Pulsed laser systems [15], [16], e.g., two-photon microscopy (2PM), can image *in-vivo* and *ex-vivo* brain tissue at micrometer scale. This approach can also be scaled by stacking *ex-vivo* slices to reconstruct whole brain anatomy with micrometer scale resolution [17], [18]. Cerebral vascular networks that can therefore be leveraged as virtual phantoms [19]. We developed a three-dimensional particle flow simulator based on highly resolved 2PM data from mice brain vasculature to conduct anatomically-realistic simulations. It was first implemented in 2D as part of a deep learning framework [20] and as part of a sparse channel sampling study for ULM [21].

In this work, we introduce a novel 3D simulation framework for the validation of ULM image formation algorithms. From a vascular model obtained using 2PM, we generate a MB flow with known MB positions in time and where dynamics mimic MB behavior from in-vivo data [7]. These MB positions at controlled concentrations are then used to position scatterers in a medium that are used to simulate ultrasound raw data, which can then be input in a ULM image formation algorithm. This setup allows to evaluate the performance of MB localization frameworks with various imaging parameters, by comparing the output MB positions with the provided reference MBs. As an example application, we studied the effect of MB concentration and transmit frequency by computing metrics such as localization error and network filling. We provide new investigations that link maximal MB concentration with the inverse of the PSF volume. Such investigations are essential in optimizing MB concentration to obtain a more complete and accurate vascular reconstruction.

## I. METHODS

### A. In-vitro two-photon microscopy

The framework starts with a highly resolved mouse brain vasculature acquired using 2PM [23] (Figure 1 A.). Prior imaging, C57BL/6 mice (25-30 g males, n = 6) were anesthetized by isoflurane (1–2% in a mixture of O2 and air) at a temperature of 37°C. The imaging apparatus consisted of a custom-built two-photon microscope with components listed in [24]. Imaging was performed through a cranial window with removed dura and a 150 μm-thick microscope coverslip for sealing purposes. Anesthesia was reduced to 0.7-1.2% isoflurane during the acquisition. Structural images of cortical vasculature were obtained by labelling blood plasma with dextran-conjugated fluorescein at 500 nM. Using angiograms acquired with a 20X Olympus objective (Numerical Aperture = 0.95), 6 stacks of vasculature (Figure 5) were prepared with voxel sizes of 1.2×1.2×2.0 μm and total volume of 600×600×662 μm.

**Figure 1:**
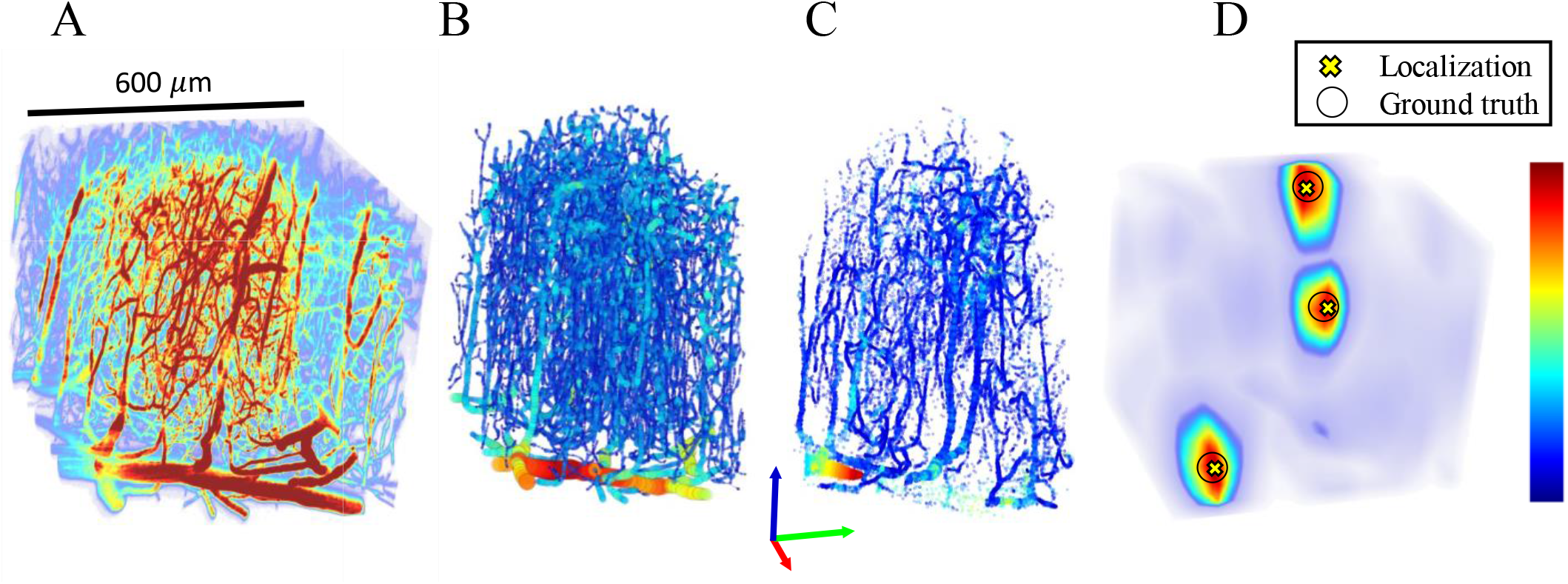
Simulation framework. All sub-figures are scaled to match the same scalebar. A) Two-photon microscopy (2PM) of in-vivo mouse brain is acquired. Colormap represents fluorescence intensity. B) From the 2PM data, a graph model is generated in 3 stages described in [22]. Colormap represents nodes’ corresponding vessel size. C) Particle trajectories are generated using the graph model and in-vivo diameter-velocity dependency. Colormap represents MB velocity. D) Using a GPU-based US simulator, RF signals are generated and reconstructed to obtain 3D+t US data, on which a correlation-based localization algorithm is applied. Colormap represents the correlation obtained using a spatial convolution of the beamformed PSF and reconstructed IQ data.

### B. Graph model

Using a modeling framework from [22], [25], a graph model of 2PM data was generated (Figure 1B). In this framework, vascular segmentation was performed using fully-convolutional neural network based on densely connected layers [22]. The graphing/skeletonization step was achieved by generating a 3D grid-graph model followed by a geometric graph contraction and refinement algorithms. Training the segmentation model was done using the Theano framework [26]. As for 3D modeling and geometric contraction processes [25], they are based on the VascGraph Python package [25]. Using this framework, a file containing vessel network geometry and topology (nodes and edges), with their corresponding inlet and outlet vessels is generated. The corresponding vessel radii are assigned as features to the nodes of the vascular network. Vessels ranged from 2 to 57.2 micrometers in diameter, their spatial position span was 500×500×650 μm and vessel volumetric density (VD) was calculated at 4%, using the ratio of vessels-containing voxels and total voxels in segmented volumes.

### C. Particle trajectory generation

Before generating a steady state flow simulation where MB concentration is always kept constant, MB trajectories were computed using the graph model source nodes, target nodes and vessels radii. For each trajectory, the shortest path between a source node and a random extremity node was calculated using vessels radii squared as weights. Since original nodes are arbitrarily spaced, a cubic spline fitting was then applied to allow for the generation MB positions with a controlled time-dependent spacing. In Hingot et al. [7], MB were followed for 5 minutes in a rat model with a continuous injection of the equivalent of 0.8 mL/kg of Sonovue MBs. MB velocities were calculated, and a log-log fit of velocity (y) as a function of vessel diameter (x) was performed on a 150 vessels sample with ln (*y*) = *p*_1_in (x) + *p*_2_ with parameters *p*_1_ = 1.9 mm μm^-1^s^-1^, *p*_2_ = −6 mm.s^-1^. Another log-log fit was computed on MB count (y) as a function of diameter (x) with parameters *p*_1_ = 3.7 MB μm^-1^, *p*_2_ = −8.2 MB, which is used in the following section (*steady-state flow*) for trajectory selection. All node-node distances (*d_i_*) were computed according to (1), where *v_i_* is the velocity from the velocity-diameter relationship of in-vivo data, d*t* is the timestamp and *P* a random normalized Poiseuille coefficient. The Poiseuille coefficient dictates the distance from the center of the vessel. The higher the coefficient, the faster is the MB and closer it is to the center of the vessel.

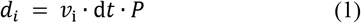

*v_i_* was calculated using the relationship with the diameter from [7]. Vessels in our network ranged from 2 to 57.2 micrometers in diameter, therefore velocities ranged from 0.01 mm/s to 5.4 mm/s. A MB trajectory is terminated if the trajectory reaches an end node or if the MB can no longer flow in the vessel due to a size constraint. In our simulations, we modelled the MBs to be of 2 micrometers in diameter, therefore all trajectories reached their maximum length. After the computation of all node-node distances (*d_i_*), the trajectory at the center of vessels (i.e., a centerline trajectory) was then used to generate other parallel trajectories to span the entire vessel. To do so, two orthogonal perpendicular vectors were calculated for each trajectory. Using a linear combination of those vectors, all orientations were created to span all possible trajectories. This step allowed for the generation of multiple trajectories at different radial positions in a specific vessel, according to their Poiseuille coefficient. Trajectories were modelled parallel to the vessel wall, i.e., being laminar as an approximation of the normal physiological state [27].

### D. Steady-state flow

Using computed microbubbles trajectories, a steady-state flow was generated using a constant concentration of MBs (Figure 1 C). The selection of microbubbles trajectories was made according to their mean diameter to match in-vivo data from [7]. To match this model, microbubble trajectories were sorted according to their mean diameter and a probability density function generated according to the model was applied for trajectories selection. To populate the steady-state flow volume, a first population of particles was generated at random positions in their respective trajectory to match a desired concentration, then, a new microbubble was added each time a microbubble reached the end of its trajectory. After this process, we were left with sets of constant number of microbubbles positions (*X*_i_,*Y*_i_,*Z*_i_) at each time frame.

### E. Scatterers’ arrangement

Unique microbubbles positions from a vascular network were randomly distributed in a 10×10×10 pattern to span over 5×5×6.5 mm (Figure 2). Since the volume of the original vascular network was minute, this setup allowed to increase the usage of the field of view and to better assess the performance of the imaging system, since the PSF is not uniform throughout the field of view. Moreover, this arrangement allowed to span a usable range of MB concentrations. For example, in the original configuration, only one MB flowing in the network corresponded to more than 5 MB/mm^3^, which is considerably high. Therefore, in this configuration, we have a 1000-fold discretization on the original MB concentration range. After ultrasound simulation, reconstruction, localization, and metrics computation, MB positions from the 1000 sub-regions were added into one original small volume to emulate a longer acquisition time only for visualization purposes (Figure 2). This step facilitated the qualitative comparison between density maps shown in Figure 7.

**Figure 2:**
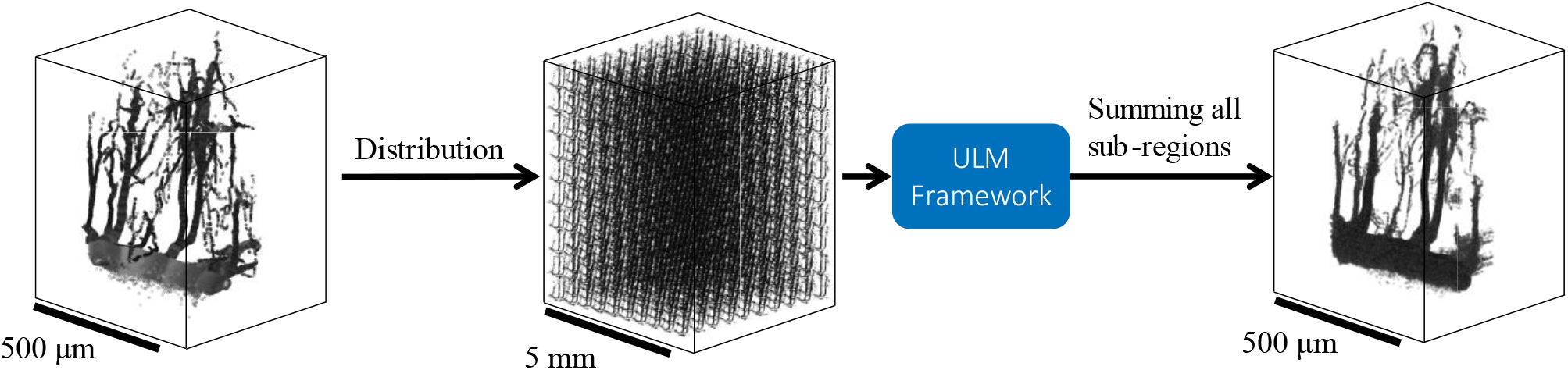
Unique microbubbles positions were randomly distributed in a 10×10×10 pattern to span over 5×5×6.5 mm. This example shows 1 million MB distributed in 1000 sub-regions, which results in 1000 MB in each sub-region. Even though fewer small vessels are present in each sub-region, this step is essential since the PSF is not uniform in the field of view and it also allows to span a usable MB concentration range. After the ULM framework and metrics computation, MB positions in sub-regions can be added back together in the original size volume to emulate a longer acquisition time for visualization purposes.

### F. Ultrasound Simulation

For 2D imaging, probe parameters matched a L22-14 linear array of 128 elements, central frequency of 15.6 MHz, pitch of 0.1 mm and element width of 0.08 mm (Vermon, France). For 3D imaging, probe parameters were set to match 32×32 matrix arrays 1024 EL 3 MHz [28], 1024 EL 8 MHz [29], [30] and the custom 1024-element 15 MHz as used by Brunner et al. [31]. The theoretical 12 MHz probe parameters were set to match the 3- and 8-MHz probes’ specifications. Bandwidths and transmit frequencies are depicted in TABLE I. Pitch and element width were set to 0.3 and 0.275 mm, respectively, the transmit pulse was a 0-degree plane wave of 4 cycles, and the effective framerate was equal to the PRF of 1 kHz.

**TABLE I.** Acquisition Parameters.

| PROBE ARRAY | TRANSMIT FREQUENCY<br>(MHz) | CENTER FREQUENCY ($C_f$)<br>(MHz) | BANDWIDTH<br>(MHz) |
| --- | --- | --- | --- |
| 128x1 | 15.625 | 15.625 | 14-22 |
| 32x32 | 3 | 3.47 | $C_f \cdot [0.7, 1.3]$ |
| 32x32 | 3.5 | 3.47 | $C_f \cdot [0.7, 1.3]$ |
| 32x32 | 6 | 7.81 | $C_f \cdot [0.7, 1.3]$ |
| 32x32 | 7 | 7.81 | $C_f \cdot [0.7, 1.3]$ |
| 32x32 | 9 | 7.81 | $C_f \cdot [0.7, 1.3]$ |
| 32x32 | 12 | 12 | $C_f \cdot [0.7, 1.3]$ |
| 32x32 | 15 | 15 | 8-22 |

To simulate the acoustic response of the microbubbles, an in house GPU implementation of the SIMUS simulation software was performed as described in [32], [33]. Gaussian white noise was added to the acoustic response to obtain a signal to noise ratio (SNR) of 15 dB in 2D and 10 dB in 3D.

### G. ULM image formation algorithm

In-phase-quadrature complex (IQ) volumes were generated from the RF data using a fast GPU-based delay-and-sum beamformer [34]. Individual MB were then identified as local maxima on correlation maps, resulting from the correlation of the reconstructed IQ volumes with the IQ spatial impulse response (PSF) of the imaging system. MB were precisely located using a 3D subpixel gaussian fitting on IQ local maxima. Localized MB were sorted as a function of their correlation from the correlation map. Only MB exceeding a certain correlation threshold were kept. Thresholding using correlation allows to screen interacting MBs. This is essential to meet the condition stated by [2], which requires isolated sources. To establish optimal threshold, tests were conducted on various concentrations at specific transmit frequencies. After finding concentration that yielded most localized MBs, localization error was plotted against correlation threshold (Figure 3). Threshold was selected to maximize MB count while preventing excessive localization error.

**Figure 3:**
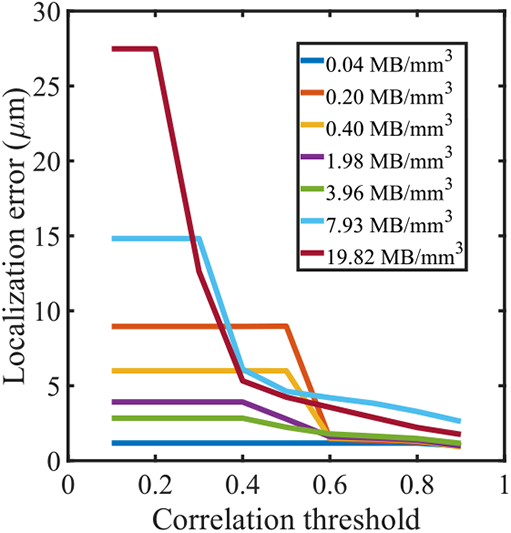
Example of a simulation conducted with the linear array at 15.625 MHz at different MB concentrations. Localization error was calculated as a function of correlation threshold. Only MB with correlation higher than the threshold were selected. In this specific case, final threshold for further simulations would be 0.6 to prevent a surge in localization error from interacting MBs.

For visualization purposes and for reconstruction times calculations, density maps were generated. MB subpixel positions were accumulated over time on a 3D sampled volume, where each voxel is approximately one fiftieth of the wavelength in size, to generate a micro vasculature density map. On a density map, each voxel intensity represents the number of microbubbles in that specific voxel.

### 2. MB simulator metrics

To validate MB behavior, 5000 MB trajectories were simulated to replicate similar MB count as in [7]. The two dependencies from *in-vivo* data are 1) the MB count as a function of vessel diameter and 2) MB velocity as a function of vessel diameter. To assess the first dependency, a histogram was computed based on the occurrences of the different diameters of all MBs at all positions. The number of histogram bins was selected as the number of integer values of the MBs radii. Natural logarithm (ln) was applied on both dependencies to generate a linear fit as in [7]. As for the second dependency, MB velocity was simply plotted against its corresponding vessel diameter.

### I. ULM metrics

To quantify the effect of concentration on the localization itself, localization error was computed using highly correlated MB positions. First, each localized MB was paired with the closest reference MB using Euclidian distance. In 2D, only in-plane distances were considered. For a specific concentration, localization error was calculated as the average distance between localized MBs and their corresponding closest reference MB. Concentration was calculated as the number of MB in a cartesian volume. This process was performed in 2D at 15.625 MHz and in 3D for different transmit frequencies spanning current usable range: 3, 3.5, 6, 7, 9 12 and 15 MHz.

Since the MB count is directly linked to reconstruction time [7], it was used as a metric to quantify the latter. After correlation thresholding, the MB count was measured at the mentioned transmit frequencies and at various concentrations. Concentrations were chosen iteratively to span a range where the MB count naturally increases with an increase in concentration and decreases when concentration is high enough that MB interactions become more important, and less MB are selected due to a lower correlation with the PSF. This allowed to find a maximum in the MB count, which was later linked to the PSF volume.

To assess effective reconstruction, density maps were generated using accumulated MB positions over different acquisition times. Selected MB concentrations corresponded to 10% of the concentrations where MB count was maximal for each transmit frequency. A reference density map with a fully populated network was generated by simulating all possible MB trajectories without a constraint on MB count (N) vs diameter. A quantitative comparison with the reference density map was made at different acquisition times using the Sørensen-Dice coefficient [35], [36]: a similarity index quantifying the network filling.

To establish the link between PSF size and MB concentration, the PSF volume was calculated for each of the different transmit frequencies and concentration at which MB count is maximal was considered as optimal. PSF was modelled as an ellipsoid with semiaxes calculated using half of the full height at maximum width (FWMH) in each dimension. For 2D imaging, ellipsoid semi-axis in the elevation plane was calculated as half of the elevation focus.

## II. RESULTS

### A. Simulated MB trajectories statistics match in-vivo data

Figure 4 displays the two dependencies that were originally extracted from in-vivo data [7]. The black lines represent the reference dependencies, colored dots represent the statistics of simulated MBs, and the blue dashed lines represent the linear fits on the simulated data. Each dot color corresponds to one of six vascular networks. Results show that the MB count increases with vessel diameter with a slope of 4.15 compared to 3.7 in reference. MB velocity also increases with vessel diameter with a slope of 1.77 compared to 1.9 in reference. Figure 5 shows the different MB flow simulations from 6 vascular networks. Each of the simulations contains an accumulated 1 million MB positions. In Figure 5, the top section depicts simulations where MB flow was constrained to follow the dependencies mentioned above. The bottom section in Figure 5 shows MB flow simulations without the constraint on MB count vs the vessel diameter. The latter simulates a random MB distribution.

**Figure 4:**
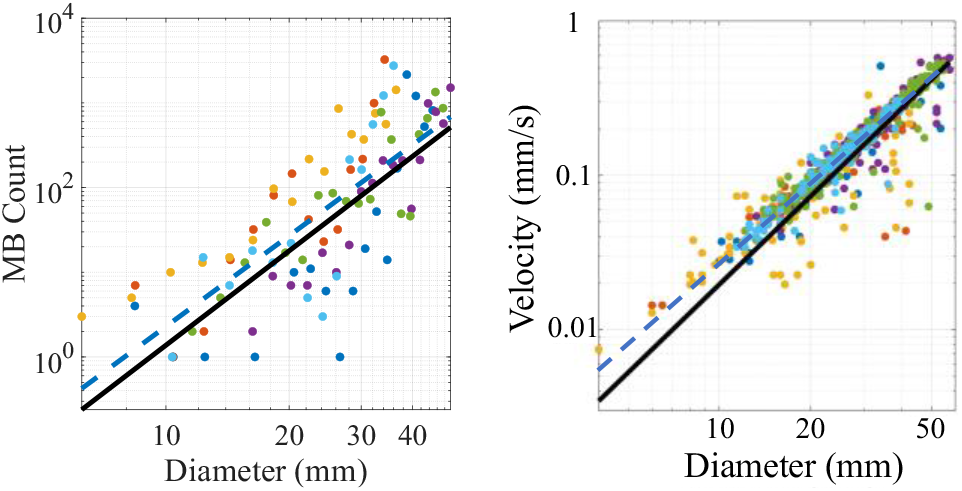
The MB flow simulator is based on two dependencies from [7], namely MB count vs vessel diameter and MB velocity vs vessel diameter. Both dependencies are in a logarithmic scale as in [7]. A number of 5000 MB trajectories were generated to match the sample statistics from [7]. (Left) A histogram of the occurrences of MB mean diameters was computed. Dots corresponds to the MB count in each histogram bin vs corresponding vessel diameter. Each dot color corresponds to one of six vascular networks. The dashed blue line represents a log-log fit as ln (y) = p_1_ln) + p_2_ with p_1_ = 3.56 and p_2_ = −7.4, R^2^ = 0.59. Diameter *x* is in mm and MB count *y* is unitless. Black line represents reference relationship with p_1_ = 3.7 and p_2_ =-8.2. (Right) Dots corresponds to MB velocities vs their corresponding vessel diameter. Each dot color corresponds to one of six vascular networks. The dashed blue line represents a log-log fit as ln(y) = p_1_ln(x) with p_1_ = 1.75 and p_2_ = −5.4, R^2^ = 0.98. Diameter *x* is in mm and velocity *y* is in mm/s. Black line represents reference relationship with pi = 1.9 and p_2_ = −6

**Figure 5:**
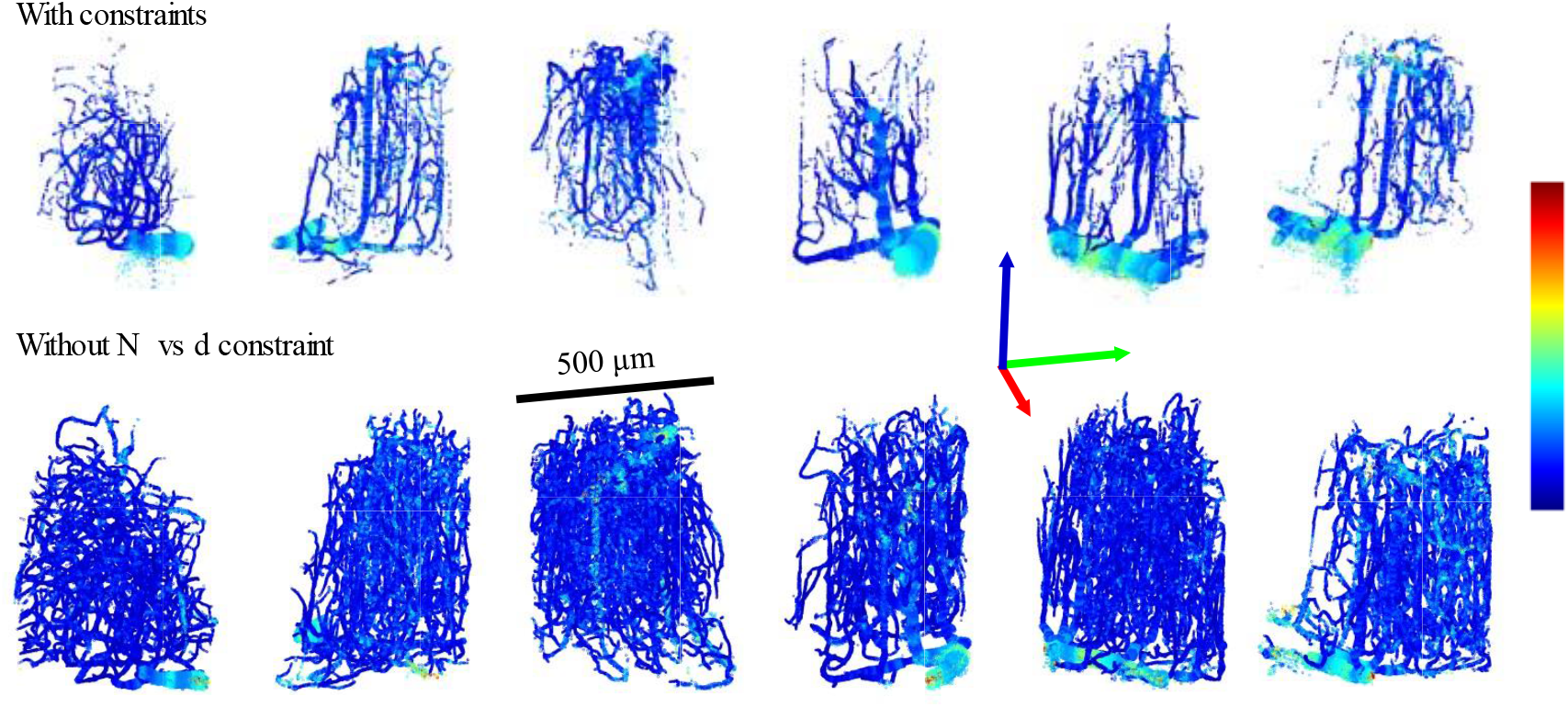
MB positions in 6 vascular networks of a mouse brain. (Top) MB flow was simulated with constraints to follow *in-vivo* dependencies: MB count and velocity with respect to the vessel diameter. (Bottom) MB flow was simulated without the MB count (N) vs vessel diameter (d) constraint.

### B. Localization error is more sensitive to concentration at lower transmit frequency

In Figure 6, localization error is depicted as a function of microbubble concentration for the different transmit frequencies. For a given transmit frequency, the bold line represents the average (μ) localization error of all localized microbubbles, and the shaded area covers the average ± the standard deviation (σ) of those localization errors. Only the in-plane distances were considered when computing 2D localization error of the linear array, i.e., the error in the elevation direction was ignored.

**Figure 6:**
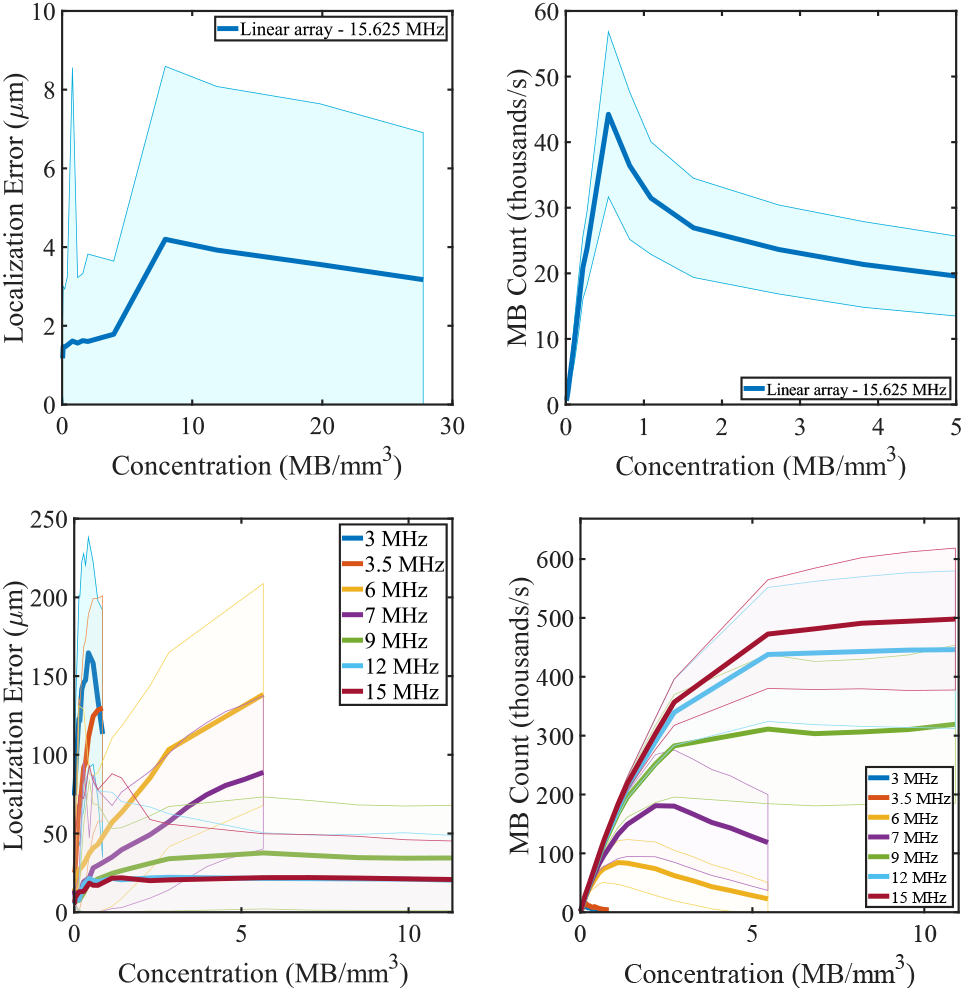
(Top left) 2D localization error is depicted as a function of microbubble concentration for the 15.625 MHz linear array. (Bottom left) 3D localization error is depicted as a function of microbubble concentration for matrix arrays at various transmit frequencies. The bold curve represents the average localization error between a localized MB and the closest reference MB. The shaded area covers the average ± standard deviation. (Top right). MB count resulting from localization as a function of concentration is depicted for the 15.625 MHz linear array. The shaded area covers the average ± standard deviation of the 6 vascular networks. (Bottom right) MB count resulting from localization as a function of concentration is depicted for different transmit frequencies using matrix arrays. Again, the shaded area covers the average ± standard deviation of the 6 vascular networks.

### C. Acquisition time is drastically reduced at higher transmit frequency

Figure 6 also shows the count of localized MB as a function of concentration. At lower frequencies, MB count increases until reaching a maximum and decreases at higher concentrations. However, from 9 MHz, MB count reaches a plateau at higher concentrations. We see that at 3-3.5 MHz, the number of detected MB is approximately 10 000 MB/s or 10 MB per frame. If we double the transmit frequency (6-7 MHz), the MB count increases with an order magnitude with approximately 100 000 to 200 000 MB/s. The rate at which the number of MB events increases seems to decrease with higher frequencies.

### D. Higher concentrations reduce reconstruction fidelity

Figure 7 shows the effect of concentration on vascular reconstruction. We applied the simulation framework in 3D using a transmit frequency of 6 MHz at four different concentrations: 0.16, 0.4, 0.8 and 1.6 MB/mm^3^. They correspond to 20, 50, 100 and 200 MB in the field of view and to 10%, 25%, 50% and 100% of the maximal MB concentration derived from Figure 6. The four concentrations were chosen iteratively, using localization error and number of localized MBs as guidelines to span a usable concentration range and to depict differences in resulting vasculature reconstruction. Top of Figure 7 shows the density maps of sub-regions of reference and localized microbubbles at increasing concentrations from left to right. Bottom of Figure 7 shows density maps acquired at different transmit frequencies, increasing from left to right. MB concentration was selected as 50% of maximal concentration (0.12 MB/mm^3^) for a 3 MHz transmit frequency.

**Figure 7:**
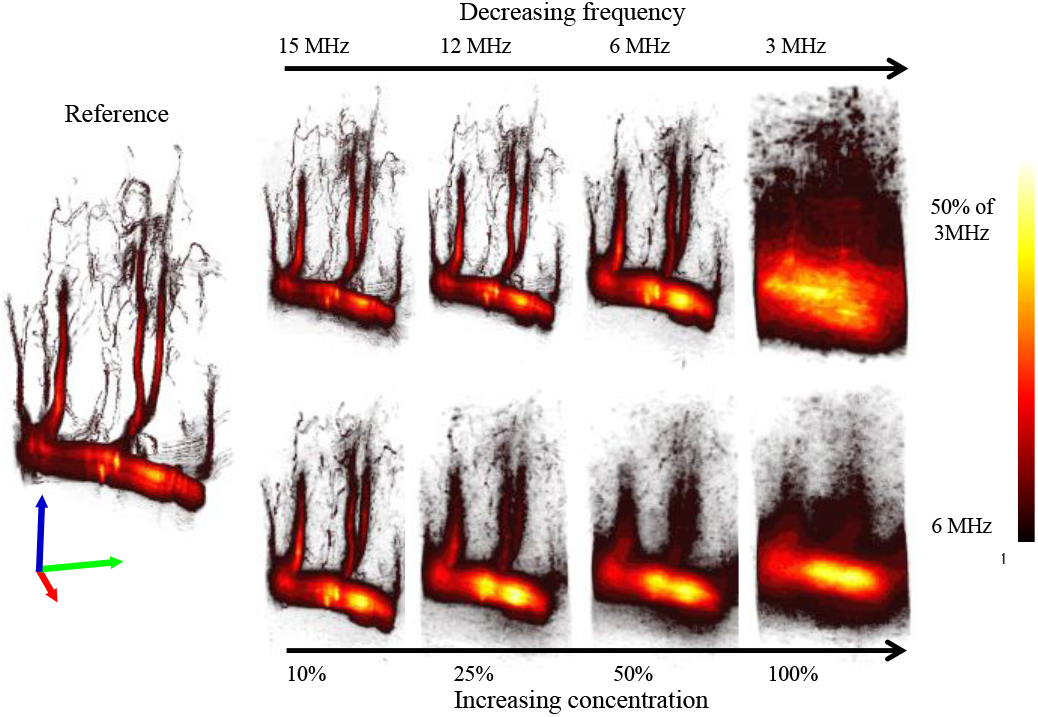
Sub-volumes of density maps of vascular network #5 corresponding to localized MB at different concentrations compared to known MB positions (ref) using a 32×32 matrix array at 6 MHz. (Top) Concentrations from low to high: 0.16, 0.4, 0.8 and 1.6 MB/mm^3^, which correspond to 20, 50, 100 and 200 MB in the field of view and to 10%, 25%, 50% and 100% of the maximal MB concentration derived from Figure 7 for a 6 MHz transmit frequency. (Bottom) For a concentration of the equivalent of 50% of the MB concentration associated with the largest number of detected MB above threshold for a 3-MHz probe (0.12 MB/mm^3^), density maps of sub-regions of simulations with transmit frequencies of 3, 6,12 and 15 MHz respectively are shown.

### E. Maximal MB concentration is PSF size dependent

Figure 8 shows the link between maximal MB concentration and PSF volume, where maximal MB concentration is proportional to the inverse of PSF volume. The uncertainty bars span the mean ± the standard deviation of the maximal MB concentration from the 6 available vascular networks.

**Figure 8:**
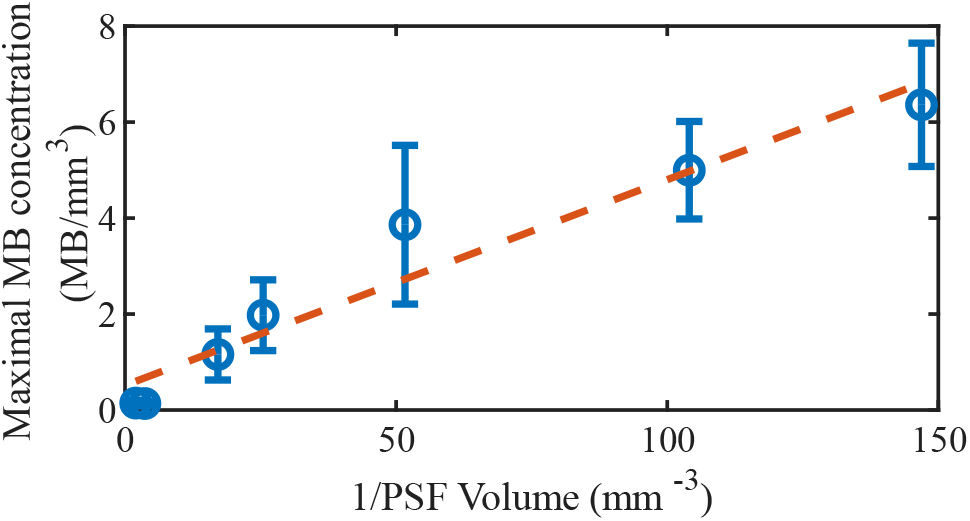
Concentration at which MB count is maximal (optimal concentration) is depicted as a function of the inverse of the beamformed PSF volume. From left to right in blue, inverse PsF volumes correspond to transmit frequencies of 3, 3.5, 6, 7, 9, 12 and 15 MHz of the matrix arrays. The dashed orange line represents a linear fit as *y* = 0.043x + 0.51, R^2^ = 0.94. The inverse PSF volume *x* is in mm^-3^ and maximal MB concentration *y* is in MB/mm^3^. The uncertainty bars span the mean ± the standard deviation of the maximal MB concentration from the 6 vascular networks.

### F. PSF volume dictates effective reconstruction time

Figure 9 shows the network filling using the Sørensen-Dice similarity index over time for the optimal concentrations corresponding to each transmit frequency.

**Figure 9:**
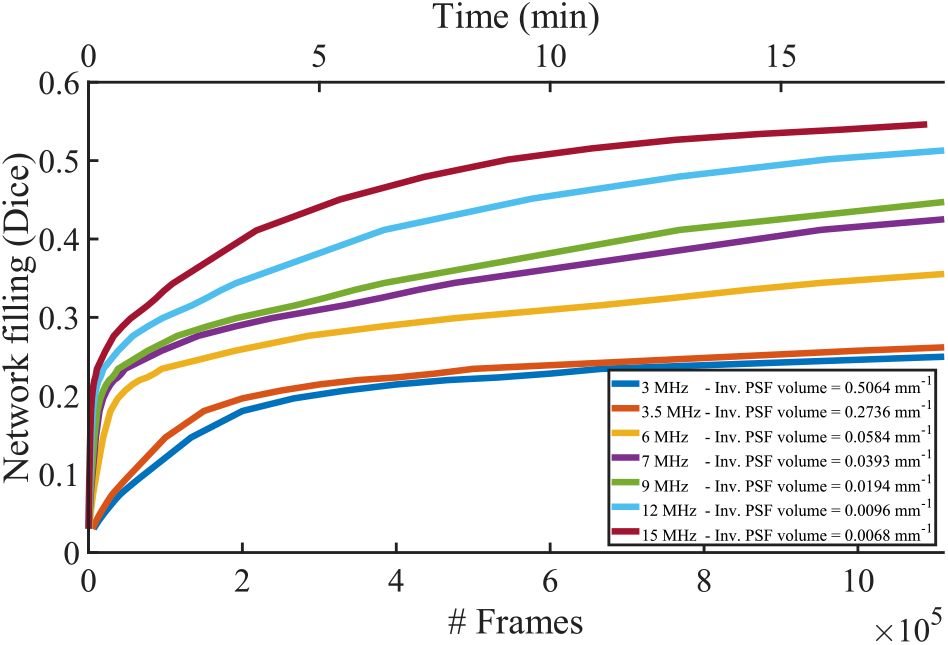
Network filling is shown using the similarity index between density maps at different acquisition times compared to a reference density map of a fully populated vascular network. MB concentrations correspond to maximal MB concentration at each transmit frequency. Time is computed considering a framerate of 1000 frames per second, where each frame corresponds to an imaging volume. MB concentration corresponds to 10% of the concentration where MB count is maximal for each transmit frequency.

## III. DISCUSSION

In this study, we took advantage of a novel 3D anatomically-realistic framework, where mice brain vasculature obtained from 2PM was used to propagate virtual MBs with velocities and MB distribution replicating behavior from in-vivo data. Having access to known MB positions at all times allowed for quantitative measurements to determine ULM capabilities as well as reference qualitative imaging. 2D and 3D localization error could be computed as well as MB count for different MB concentrations and transmit frequencies. A link between maximal MB concentration and PSF volume was found and network filling as a function of time was calculated. Results allow to answer fundamental questions related to in-vivo imaging such as whether the vasculature is accurately or fully depicted as imaging parameters such as the probe or concentrations used are varied.

Simulations were conducted on the 6 vascular networks separately. The variety of vascular morphologies allowed to generalize the link between MB concentration and acquisition parameters while reducing potential overfitting.

### A. MB simulation

Figure 4 shows that our MB simulation framework was successful at replicating observed MB behavior from in-vivo dependencies, namely the MB count and velocity vs vessel diameter. Due to these dependencies, realistic MB flow and MB distribution are possible in subsequent ultrasound simulations.

Figure 5 shows that constraints on MB flow favor MB distribution towards larger vessels. Visually, fewer small vessels are mapped for a set amount of MBs, but large vessels appear fuller. This means that for a set acquisition time and a set MB concentration, some small vessels will not or will not be fully mapped in the final density map regardless of imaging resolution.

### B. MB localization

In Figure 7, we see that as concentration increases, even though most vessels are preserved, they appear blurred, are missing, or are inaccurately depicted from the reference density map. This is in accordance with theory since ULM requires isolated MB signals. The higher the concentration, the more likely MBs will interact with each other. However, relative concentration around 10% of the concentration at maximal MB count allows to depict most of the original vasculature. Furthermore, reference density map shows that because of realistic MB count vs diameter, some small vessels are not present. Probability of a MB flowing in vessels of only a few micrometers is so low that it would require a longer acquisition time than the equivalent of 10 minutes at 15 MHz, according to results in Figure 9. This means that in current in-vivo imaging, some vessels of only a few micrometers are most likely not mapped if the transmit frequency is not high enough or the acquisition time not long enough.

In Figure 6, the decrease in MB count at high MB concentrations shows that thresholding MBs using their correlation with beamformed PSF allows to screen for intensely interacting MBs. Localization error increases with concentration as predicted by theory and is in conjunction with vessels blurring or missing. Localization error can reach a plateau at higher frequencies when selected MBs becomes constant due to correlation thresholding. Higher transmit frequencies are associated with lower localization errors as predicted by theory [37]. Higher frequencies are associated with smaller PSFs; hence a higher concentration is possible while maintaining isolated sources.

In Figure 6, the rate at which MB count increases eventually slows down with higher frequencies. This is mainly due to the size of the PSF only decreasing in the axial direction because of a constant element pitch and width. This is due to current manufacturing capabilities that limit piezoelectric element size.

The dependency in Figure 8 shows that PSF volume can be used predict maximal MB concentration. In theory, to reduce MB interactions, MB concentration must be decreased. We see here that reducing PSF size also reduces MB interactions. Since MB concentration should be as high as possible to reduce acquisition time, this dependency can be used for choosing optimal concentration for a specific probe’s acquisition parameters.

In Figure 9, the network filling represents the intersection between reference density maps and density maps resulting from MB localization at different transmit frequencies. The higher the network filling index, the more of reference vessels are present in the mapped vasculature from the imaging setup. Results in Figure 9 show that acquisition time is drastically reduced using smaller PSF volumes (or higher inverse PSF volumes). This means that a higher transmit frequency should be prioritized over a lower transmit frequency if the required penetration depth allows for it. Also, for a set acquisition time, depending on MB concentration and PSF volume, only a portion of total vasculature is mapped.

### C. Limitations

This study set forth some limitations of the proposed simulation framework.

- A graph model was generated from 2PM, through automated segmentation, surface modeling and contraction. From that, MB trajectories were generated using MB velocities and number of MB as a function of diameter from two models derived from in-vivo data. This approach guarantees anatomically-realistic modelling since vasculature is derived from an ex-vivo model. Another avenue would consist of generating a fully synthetic fluid flow-driven graph model as in [38], where vasculature is generated using constraining parameters, namely vascular resistance, diameter, pressure, and bifurcation position. Whole cortical circulation can be generated while distinguishing between arterial and venal flow. This framework combines anatomical data with artificial construction laws to overcome limitations in coverage and resolution in reference anatomical data.
- In ultrasound localization microscopy, there are many parameters that can influence SNR, contrast, resolution, network filling and so forth, hence influencing the accuracy of the vascular reconstruction. In this study, we chose parameters that are most closely linked to our ability to maintain isolated sources in a spatiotemporal referential as stated in [2]. For that reason, we focused primarily on MB concentration [7] and transmit frequency [6].
- In Figure 4, variability in the MB count vs vessel diameter is due to the specificity of the vascular network. The smaller the vascular network, the more significant this variability is. Variability in MB velocities vs vessel diameter can be due to abrupt vessel diameter change due to rough vascular segmentation, which can affect the accuracy of a MB’s corresponding vessel diameter prediction.
- As for ultrasound signals, they were generated using either readily available or realistic ultrasound probes acquisition parameters. The rationale behind this was to assess current limitations and seek for optimal ULM parameters. White Gaussian noise was added in radio frequency data to improve realism, but no tissue was modeled. Since PSF size and shape greatly influences our ability to precisely detect single MB events, any alteration to the PSF from higher noise or lower contrast from neighboring tissue could have an impact on MB localization and ultimately maximal MB concentration. Other studies with tissue modelling are therefore to be conducted with tissue modelling as in [39], [40].
- In Figure 6, we can see that thresholding does not completely prevent MB interactions, as depicted in the increasing localization error as a function of concentration. Therefore, correlation threshold must be selected wisely to maximize MB detection events for a fuller network while minimizing strongly interreacting MBs which is depicted as blurred vasculature in density maps.
- All simulations were conducted on networks of brain mice vasculature where MB flow reflected behavior from in-vivo data. However, the vascular networks were relatively small in terms of field of view. This means that MB trajectories are limited to a small range of vessel sizes (mainly small vessels). This reduced vessel size range increases inter-network variability which can be seen through results from Figure 8, where MB count variability is not proportional to the inverse of PSF volume. In the medium, MBs were spread uniformly to artificially cover a large portion of space, but realism could be improved by having access to a larger network and ultimately a whole brain or whole organ vasculature, where the vascular network is closer to real-life vascular size distribution, which will increase metrics repeatability.
- In subsequent studies, acquisition parameters such as number of compound angles [34], number of pulse cycles, pulse frequency modulation (chirp) [41], SNR, probe elements geometry, MB size distribution just to name a few could be interesting to better understand the underlying limitations of ULM and progress towards finding optimal imaging parameters.

## IV. CONCLUSION

In conclusion, we have developed a framework capable of characterizing ULM image formation algorithms. We described how to obtain virtual anatomically-realistic MB positions from 2PM data. We have shown and quantified the impact of concentration on MB localization and acquisition time. We quantified the link between maximal MB concentration and PSF volume. Standardized anatomically-realistic MB simulations could become a useful tool in the validation of an ULM imaging setup and the PSF volume a straightforward metric for a specific probe’s acquisition parameters since it would define the appropriate MB concentration. In practice, we hope that these results can be used to achieve optimal vasculature reconstruction at lowest acquisition time for a specific imaging setup.

